# Tissue-specific mRNA profiling of the *Brassica napus-Sclerotinia sclerotiorum* interaction uncovers novel regulators of plant immunity

**DOI:** 10.1101/2021.03.27.437327

**Authors:** Philip L. Walker, Ian J. Girard, Shayna Giesbrecht, Steve Whyard, W.G. Dilantha Fernando, Teresa R. de Kievit, Mark F. Belmonte

## Abstract

White mold in *Brassica napus* (canola) is caused by the fungal pathogen *Sclerotinia sclerotiorum* and is responsible for significant losses in crop yield across the globe. With advances in high-throughput transcriptomics, our understanding of the *B. napus* defense response to *S. sclerotiorum* is becoming clearer; however, the response of individual tissue layers directly at the site of infection has yet to be explored. Using laser microdissection coupled with RNA sequencing, we profiled the epidermis, mesophyll and vascular leaf tissue layers in response to *S. sclerotiorum*. This strategy increases the number of genes detected compared to whole-leaf assessments and provides unprecedented information on tissue-specific gene expression networks in response to pathogen attack. Our findings provide novel insight into the conserved and specific roles of ontogenetically distinct leaf tissue layers in response to infection. Using bioinformatics tools, we identified several defense genes that might coordinate plant immunity responses shared across different tissue layers within the leaf. These genes were functionally characterized by challenging T-DNA insertion lines of Arabidopsis with necrotrophic, hemi-biotrophic, and biotrophic pathogens, ultimately converging on the PR5-like RECEPTOR KINASE (PRK5). Together, these data provide insight on the complexity of the *B. napus* defense response directly at the site of infection.

**Highlight:** Laser microdissection coupled RNA sequencing of the *B. napus* – *S. sclerotiorum* interaction identifies candidate genes predicted to guide plant immunity against pathogen attack.

## Introduction

*Brassica napus* L. (canola, oilseed rape) is a global oilseed crop and is susceptible to numerous fungal and bacterial pathogens (Neik et al., 2017). Response of *B. napus* to pathogen attack involves multiple immunity and induced resistance pathways which can be differentially activated depending on whether the pathogen has a necrotrophic, hemi-biotrophic, or biotrophic lifestyle. Systemic acquired resistance (SAR) and induced systemic resistance (ISR) pathways are activated in response to infection to induce physical and chemical defense mechanisms locally and throughout the host plant (Corné et al., 2014). These pathways are mediated by the expression of defense hormones; SAR being mediated by salicylic acid (SA), and ISR being mediated by jasmonic acid (JA) and ethylene (ET) (Yuan et al., 2019). JA/ET mediated ISR has been associated with necrotrophic infection, while SA and SAR are commonly associated with response to biotrophic infection (Shen et al., 2018). Further, induction of a local hypersensitive response in SAR can be used opportunistically by necrotrophic pathogens that feed on dead cells (Shen et al., 2018). However, the relationships between these defense hormones and the inherent crosstalk between resistance pathways is still unclear as they share common regulators whose activation can lead to antagonistic expression (Robert-Seilaniantz et al., 2011), or in some cases, co-expression leading to synergistic effects and a stronger immune response to pathogen attack (Ngou et al., 2020). Two of the plant immunity pathways, pattern-triggered immunity (PTI) and effector-triggered immunity (ETI), function in pathogen detection and initiation of host defense at the site of infection, and like resistance pathways, have shown specific and shared modes of activation and associated downstream defense processes. PTI has previously been implicated in successful defense against broad-spectrum necrotrophs (Mengiste 2012; Wang et al., 2018) and is characterized by receptor-like kinases (RLKs) acting as pattern recognition receptors (PRRs) capable of detecting pathogen associated molecule patterns (PAMPs) (Saijo et al., 2017; Tang et al., 2017). This leads to downstream signal transduction cascades and the up-regulation of defense hormones and genes associated with an oxidative burst within the cell, callose deposition, phytoalexin production, and cell wall fortification (Tang et al., 2017; Henry et al., 2014; Nasir et al., 2018). In contrast, ETI is commonly associated with nucleotide-binding site leucine-rich repeats (NBS-LRR) pathogen detection and response to biotrophic pathogens, and like SAR, is characterized by the activation of a local hypersensitive response at the site of infection (Ding et al., 2011). However, similar to the relationship of SAR and ISR pathways, PTI and ETI immunity pathways have shown potential synergistic effects when co-induced, leading to increased pathogen resistance (Zhang et al., 2018; Henry et al., 2014). While these pathways have been well studied at the whole leaf level in response to pathogen attack (Girard et al., 2017; Chittem et al., 2020), the activation of these pathways and down-stream defense responses at the tissue-specific level has yet to be explored.

*Sclerotinia sclerotiorum* (Lib) de Bary (del Rio et al., 2007), the causal agent of white mold, contributes to significant yield losses in *B. napus* and several other important crops species (Wang et al., 2019). *S. sclerotiorum* activates a suite of cutinases and lipases to penetrate the leaf cuticle, as well as the pathogenicity factor oxalic acid which functions in modulating the host redox environment (Seifbarghi et al., 2017). This is followed by the release of numerous hydrolytic and cell wall degrading enzymes to break down cell wall-associated polysaccharides (Seifbarghi et al., 2017; Bashi et al., 2012). *S. sclerotiorum* is an aggressive necrotrophic pathogen that penetrates the epidermis and mesophyll tissues of *B. napus* within 24 hours post inoculation (hpi) and colonizes all leaf tissues including the vasculature within 48 hours (Girard et al., 2017; Uloth et al., 2015). In response to necrotrophic infection, whole-leaf RNA sequencing revealed enrichment of the JA/ET associated PTI and ISR pathway (Girard et al., 2017; Joshi et al., 2016). Tolerance to the pathogen was shown to operate through redox homeostasis and the activation of PTI/ISR associated ethylene response factors, thereby limiting host damage by pathogen induced reactive oxygen species (ROS) production (Girard et al., 2017).

To better understand how *B. napus* responds to *S. sclerotiorum* directly at the site of infection, we profiled tissue-specific and shared RNA populations in the epidermis, mesophyll, and vascular tissues 24 hpi. The transcriptomic analyses revealed novel tissue-specific and shared mRNAs which were not identified previously through whole-leaf sequencing (Girard et al, 2017). Functional characterization of selected candidate genes in Arabidopsis T-DNA insertion lines identified multiple plant immunity signalling molecules and transcriptional regulators that play an essential role in successful pathogen defense following inoculation with necrotrophic, hemi-biotrophic, and biotrophic fungal pathogens. Taken together, we have identified both tissue-specific and shared immunity pathways involved in both PTI and ETI and investigate putative mechanisms responsible for plant-pathogen interactions.

## Materials and methods

### Plant growth & *S. sclerotiorum* inoculation

*B. napus* seeds were stored and grown according to Girard et al., 2017. *Sclerotinia sclerotiorum* ascospores were collected at the Agriculture and Agri-Food Canada Morden Research and Developmental Centre (Morden, Manitoba, Canada) and stored on aluminum foil at 4°C. Ascospore inoculum was prepared by suspending 8×10^4^ spores/ml in a 15 ml distilled water and 0.01% Tween20 solution. *B. napus* petals were inoculated with 30 µl of ascospore inoculum and stored at room temperature for 72 hours in a Parafilm sealed petri plate. Control petals were inoculated with a 0.01% Tween20 solution and stored under the same conditions. *S. sclerotiorum* infected and control petals were transferred to the adaxial side of canola leaves at the 30 – 50% bloom stage. Inoculated leaves were covered using plastic bags to increase humidity levels for optimal *S. sclerotiorum* pathogenicity. Infection sites were subsequently collected at 24 hpi for tissue processing and laser microdissection.

### Tissue processing & laser-capture microdissection

Infected canola leaves were collected and processed for LMD as described in Belmonte *et al*. (2013). Infection sites were collected from leaf tissue and fixed in 3:1 (v/v) ethanol:acetic acid overnight at 4°C. Tissues were subsequently rinsed and dehydrated in a graded ethanol series (75%, 85%, 95%, 100%, 100%) followed by xylene infiltration (3:1, 1:1, 1:3 ethanol:xylene (v/v), 100% xylenes, 100% xylenes) at 4°C overnight. 100% xylene and paraffin wax chips were then added to the xylene-infiltrated tissue prior to being stored at 4°C overnight. Tissues in xylene were then brought to room temperature (∼21°C) and incubated at 38°C for 60 minutes followed by 57°C for 30 minutes. Five changes of 100% paraffin were performed one hour apart prior to embedding. Processed leaf tissues were then sectioned using a Leica RM2125RT rotary microtome at 10 μm under RNase-free conditions and mounted on Leica PEN membrane slides before being deparaffinized in xylene twice for 30 seconds per wash. Histological sections were collected using the Leica Laser Microdissection 7000 system in lysis buffer (Ambion, Austin, TX, USA) for downstream RNA isolation. Epidermis, mesophyll, and vascular tissue layers were collected separately with each bio-replicate being collected from at least 30 different plants. Each tissue layer was collected independently of each other with infected samples being collected in a 400 µm area directly adjacent to the infection site to ensure cells collected were actively responding to infection and at a similar stage of response for each collection (Fig. 1a).

**Figure 1.**
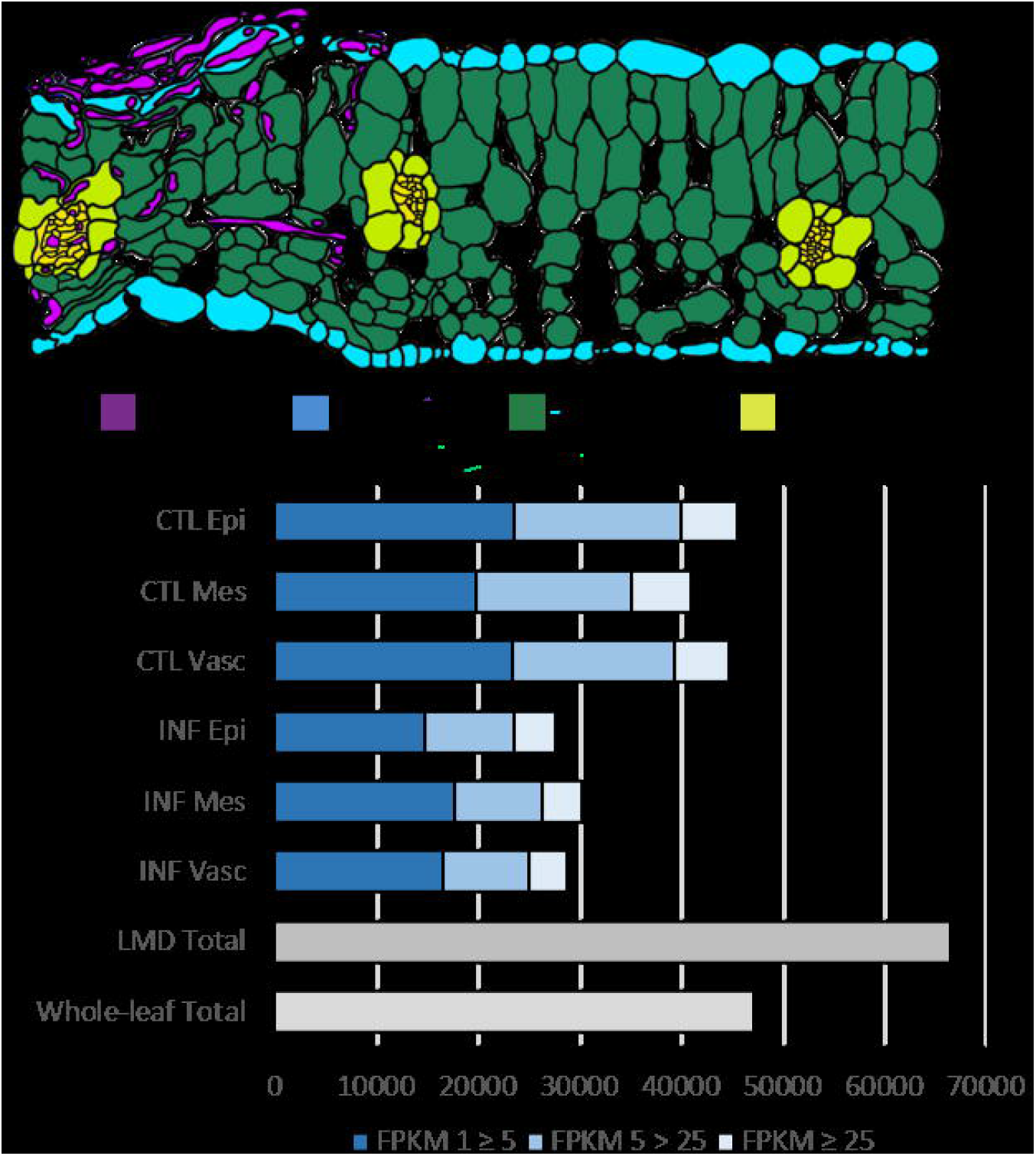
Diagram of an infected *B. napus* leaf cross-section and global gene activity in the *B. napus-S. sclerotiorum* pathosystem. A) *S. sclerotiorum* infected mature *B. napus* leaf prior to collection of individual tissue layers under laser-capture microdissection. Collection area of 400 µm adjacent to the infection site for infected tissue. B) Hierarchical clustering and global gene activity of epidermis, mesophyll, and vascular leaf tissue in *B. napus*. Number of transcripts detected across all treatments were identified through RNA sequencing. Transcripts with an FPKM [Fragments Per Kilobase of transcript per Million mapped reads] > 1 are considered to be detected. Detected transcripts are subdivided into low (FPKM 1 ≥ 5), moderate (FPKM 5 > 25), or high (FPKM ≥ 25) abundance levels. CTL = control, INF = infected, LMD = laser microdissection.

### cDNA library preparation & RNA sequencing

RNA was isolated using the Ambion^®^ RNaqueous^®^ micro kit according to manufacturer’s instructions. RNA quality was analyzed using the Agilent 6000 Pico LabChip^®^ and Agilent 2100 Bioanalyzer software (Agilent Technologies, USA). RNA quantity was determined using the Quant-iT™ RiboGreen^®^ kit according to manufacturer’s instructions. cDNA library synthesis was performed according to Ziegler et al. (2019). Subsequently, 100 bp single-end RNA sequencing was performed using the Illumina HiSeq2000 platform (Génome Québec Innovation Centre, McGill University, Montreal, Canada).

### Data analysis of RNA sequencing data

Raw sequencing reads were deposited at GEO (GSE169299). Read quality control and adapter sequence removal was performed using Trimmomatic 0.36 (Bolger et al. 2014) (HEADCROP:9 LEADING:30 TRAILING:30 SLIDINGWINDOW:4:30 MINLEN:50). Surviving reads were subsequently aligned to the *B. napus* genome v4.1 (Chalhoub et al. 2014) using HISAT2 (Kim et al. 2015) alignment software. The cufflinks suite (Trapnell et al. 2012) was used to generate normalized read counts in FPKM (fragments per kilobase of transcript per million mapped reads). FPKM expression tables can be found in Dataset S1.

### Hierarchical and fuzzy k-means clustering

Hierarchical and fuzzy k-means (FKM) clustering was performed using the pvclust (https://cran.r-project.org/web/packages/pvclust/index.html) and cluster http://cran.r-project.org/web/packages/cluster/cluster.pdf) packages respectively in R studio. Average FPKM values for all treatments were used as input for both clustering analyses, with a detection cut off ≥ 1 FPKM. FKM pattern gene lists can be found in Dataset S2.

### Gene ontology (GO) term enrichment

SeqEnrich (Becker et al. 2017) was used to perform GO enrichment analysis on FKM gene lists. GO terms were considered significantly enriched at P ≤ 0.001 or a log^10^ p-value ≤ −3. GO summary tables can be found in Dataset S3. GO heatmap visualization was performed using the conditional formatting function in Excel.

### T-DNA insertion Arabidopsis susceptibility screening against multiple pathogens

To screen for potential susceptibility phenotypes, nine Arabidopsis Col-0 loss of function T-DNA insertion lines were challenged with fungal and bacterial pathogens. T-DNA mutants were obtained from the Arabidopsis Biological Resource Centre (https://abrc.osu.edu/) and confirmed for homozygous T-DNA insertions using PCR. Arabidopsis seeds were sterilized using a graded ethanol series (70%, 70%, 95%) before being plated on MS media for a 3-day vernalization at 4°C. Seeds were then incubated at 22°C under constant light for 10 days. Seedlings were transplanted in Sunshine Mix #1 soil and grown under long day (16 hours light, 8 hours dark 150-200μE/*m*^2^ /s) conditions at 22°C. Arabidopsis seedlings matured for 14 days prior to inoculation and a minimum of n = 15 leaves were inoculated per treatment. Two leaves per bio-replicate and 3 bio-replicates were collected for qPCR fungal load quantification post-inoculation, described below.

### Bacterial and fungal infection and quantification assays

Ascospore inoculum for both *S. sclerotiorum* and *B. cinerea* contained 1×10^6^ spores/ml suspended in 10 ml potato dextrose broth + peptone solution and 0.015% Silwet L-77. 10 µl of inoculum was pipetted onto the midvein of mature Arabidopsis leaves and stored in sealed humidity chambers to allow for optimal infection conditions (Mcloughlin et al., 2018). Leaves were subsequently imaged and collected at 3 days post inoculation (dpi). Lesion size in mm^2^ was quantified using ImageJ software (imagej.nih.gov).

*L. maculans* conidiospore inoculum contained 1×10^7^ spores/ml in sterile water. Leaves were point inoculated according to Becker et al. (2017) using 10 µl of conidiospore inoculum and subsequently imaged and collected at 20 dpi.

*H. arabidopsidis* infection assays were performed using fresh spray inoculum of *H. arabidopsisidis* isolate Emco5 conidiospores according to Shirano et al., 2002. Arabidopsis seedling lawns were grown under long day conditions at 18°C and inoculated at 10 days maturity. Seedlings were imaged using the Leica MZ6 dissecting microscope at 14dpi and subsequently collected for fungal load qPCR analysis

Fungal load was quantified using qPCR of 18S rRNA of *S. sclerotiorum, B. cinerea* in both mutant and col-0 Arabidopsis, while actin was used for *L. maculans* and *H. arabidopsidis* quantification. Each treatment group consisted of 3 bio-replicates, with each bio-replicate containing at least three infected leaves. The ΔCt method was used to analyze mRNA abundance in mutants relative to Col-0 control using infected mature Arabidopsis leaves as input tissue. Collection timepoints were selected for each pathogen tested based on how aggressive infection progressed, not allowing collection timepoints to exceed 100% lesion area in susceptibility phenotypes. A list of primers used to detect fungal load is found in Dataset S4.

*P. syringae* pv. *tomato* DC3000 spray inoculum was prepared and applied according to Laflamme et al. (2016). Infected leaves were imaged and collected at 5 dpi for subsequent colony plate quantification. Infected leaf tissue was ground in 10 mM MgSO4 prior to being plated on Pseudomonas isolation agar containing rifampicin and cycloheximide. Plates were incubated at 28°C for 2 days before colony quantification.

### Analysis of PR5K predictive interacting partners

Predictive PR5K interacting partners were identified using stringdb (https://string-db.org/) and co-expression data. Gene candidates were quantified for their expression levels in *prk5* mutants compared to Col-0 at 3 dpi in the presence of *S. sclerotiorum*. qPCR analysis was performed using three biological replicates per treatment and transcript abundance calculated using the ΔΔCT method with EF1BB (AT1G30230) as a housekeeping gene. A list of primers used to study predictive PR5K interacting partners is found in Dataset S4.

## Results

### LMD-RNAseq increases spatial resolution of mRNA populations of *B. napus* leaf tissues in response to *S. sclerotiorum*

We profiled the tissue-specific defense response in *B. napus* leaf tissues infected with *S. sclerotiorum* using a combination of LMD and RNA sequencing (Fig. 1a). Hierarchical clustering of FPKM (fragments per kilobase of transcript per million mapped reads) expression values reveal distinct patterns of differential transcript abundance between control and infected tissues (Fig. 1b). Data show two distinct clusters based on fungal treatment. Within the control and infected clusters, leaf epidermal and mesophyll tissue layers grouped together and demonstrated distinct expression patterns compared to vascular tissue (Fig. 1b). The union of all mRNA populations in both control and infected treatment groups identified a total of 66 454 transcripts with an abundance value ≥ 1 FPKM, compared to the 47 154 transcripts detected previously using whole-leaf sequencing (Girard et al. 2017). Increased transcript detection was found in control versus infected tissue with an average of 43 758 and 28 884 transcripts per treatment identified respectively and is likely due to increased *S. sclerotiorum* transcript detection in infected samples (Dataset S5). However, both control and infected treatments demonstrated similar levels of detected transcripts. For example, we detected an average of 25% high (FPKM ≥ 25), 35% moderate (FPKM 5 > 25), and 50% low abundance (FPKM 1 ≥ 5) transcripts in control tissues, while infected tissue identified 24% high, 29% moderate, and 57% low abundance mRNAs (Fig. 1b).

### Fuzzy k-means clustering analysis reveals tissue-specific biological processes in response to *S. sclerotiorum* infection

Next, we identified tissue-specific and shared defense gene expression patterns in *B. napus* leaves using fuzzy k-means (FKM) clustering. Fig. 2A shows a co-expression pattern of genes shared across all three tissue systems of the leaf (2 040 genes) that are up-regulated in response to *S. sclerotiorum*. We also find tissue-specific patterns in response to infection, with an epidermis-specific pattern of 128 genes, mesophyll-specific pattern of 394 genes, and a vascular-specific pattern of 267 genes. To identify biological processes and molecular functions associated with these patterns, we performed a gene ontology (GO) term enrichment analysis on genes belonging to these co-expressed gene sets (Fig. 2b). The shared pattern showed enrichment of terms associated with multiple facets of the defense response including pathogen detection and signal transduction (chitinase activity P = 2.56×10^−11^, ser/thr kinase P = 1.50×10^−6^, pattern recognition receptor activity P = 4.09×10^−4^, and chitosan binding P = 4.09×10^−4^), plant defense hormones (salicylic acid P = 2.0×10^−10^, jasmonic acid P = 5.94×10^−5^, brassinosteroid P = 1.87×10^−4^ signalling pathways and ethylene biosynthesis P = 5.01×10^−4^), vesicle-mediated exocytosis (SNARE complex P = 1.90×10^−6^, exocytosis P = 1.18×10^−4^), cell wall reinforcement (callose deposition P = 1.65×10^−9^, pectin biosynthesis P = 7.08×10^−4^, cell wall biogenesis P = 3.05×10^−4^), and the respiratory burst (response to hypoxia P = 5.64×10^−12^, respiratory burst P = 4.34×10^−9^, glutathione biosynthesis P = 2.73×10^−6^, and response to reactive oxygen species P = 2.67×10^−4^). The epidermis-specific pattern showed GO terms associated with ethylene-activated signalling (P = 1.02×10^−6^) and polyamine biosynthesis (P = 7.36×10^−5^) pathways. The importance of these molecules in the epidermis was supported by the co-enrichment of adenosylmethionine decarboxylase activity (P = 7.84×10^−6^) and is involved in both ethylene and polyamine biosynthesis (Majumdar et al., 2017). Further, we identified enrichment of cell redox homeostasis (P = 7.14×10^−5^), a key function of polyamines in plants (Rider et al., 2007). The mesophyll tissue showed a strong association with GO terms related to structural reinforcement in the cell wall, including peroxidase activity (P = 4.87×10^−31^), lignin biosynthesis (P = 6.28×10^−6^), xyloglucan metabolism (P = 1.46×10^−6^), and cell wall organization (P = 1.70×10^−18^). Lastly, the vascular tissue was enriched for GO terms involved in induced resistance pathways, SAR (response to salicylic acid P = 4.32×10^−12^, and SAR-involved salicylic acid mediated pathway P = 3.73×10^−4^) and ISR (ISR-involved jasmonic acid mediated pathway P = 6.33×10^−7^, ethylene-activated signalling pathway P = 1.77×10^−4^, and oxylipin biosynthesis P = 7.60×10^−6^). We also find the regulation of cell death (P = 6.93×10^−6^), leaf senescence (P = 2.34×10^−7^), and G-protein coupled receptor signalling (P = 7.55×10^−6^) suggesting a role in programmed cell death (PCD) in the vascular tissue. Taken together, data suggests cells and tissues of the leaf respond both specifically and coordinately in response to *S. sclerotiorum* infection.

**Figure 2.**
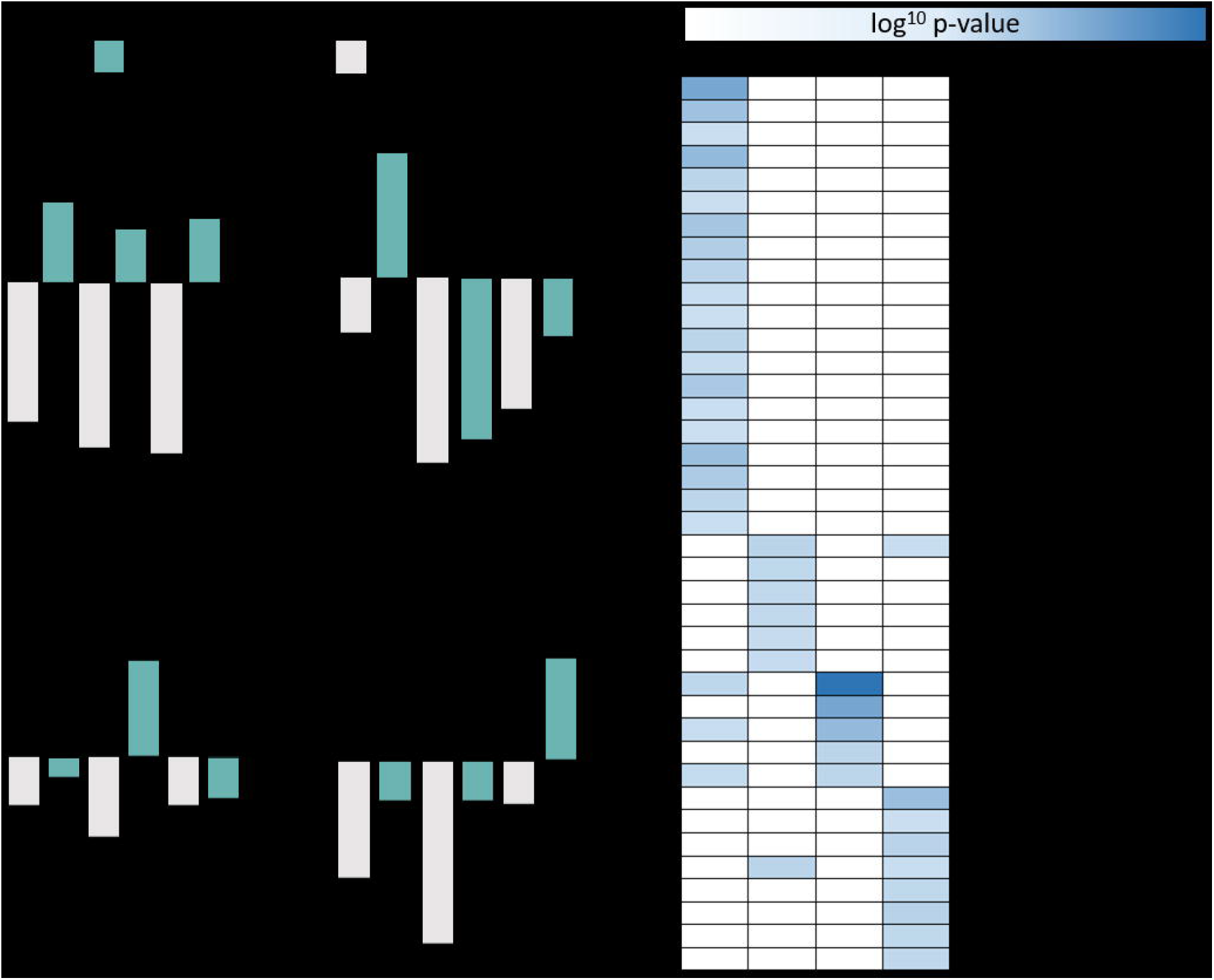
Identification of tissue-specific and shared patterns in response to *S. sclerotiorum* infection using fuzzy k-means clustering and GO enrichment. A) Fuzzy k-means clustering bar plots of mapped genes demonstrating an FPKM abundance value ≥ 1. B) Heatmap of enriched GO terms associated with shared and tissue-specific up-regulated gene patterns. GO terms considered statistically significant with a hypergeometric p-value < 0.001.

### Functional characterization of defense-related genes against fungal and bacterial pathogens using Arabidopsis T-DNA insertion lines

Next, we challenged Arabidopsis T-DNA insertion lines that showed a reduction in defense gene activity with *S. sclerotiorum* and other fungal and bacterial pathogens to explore potential susceptibility phenotypes. A total of nine genes were selected based on their presence in FKM clustering patterns as well as their up-regulation in *S. sclerotiorum*-infected tissue and associated processes (Table 1). Four of the selected genes demonstrate tissue-specific up-regulation (*AT4G39940, AT1G25340, AT3G25882, AT1G59870*) and five demonstrate shared up-regulation across all tissue layers (*AT4G18250, AT1G76470, AT5G41750, AT5G38340, AT3G11820*). The Arabidopsis mutant lines and wild type (Col-0) plants were challenged with the closely related necrotrophic fungal pathogens *S. sclerotiorum* and *Botrytis cinerea;* the hemi-biotrophic fungal pathogen *Leptospahaeria maculans*; the biotrophic fungal pathogen *Hyalonopernospora arabidopsidis*, and the bacterial biotrophic pathogen *Pseudomonas syringae*. These pathogens were selected to explore defense roles against pathogens with diverse pathogenic lifestyles.

**Table 1.**
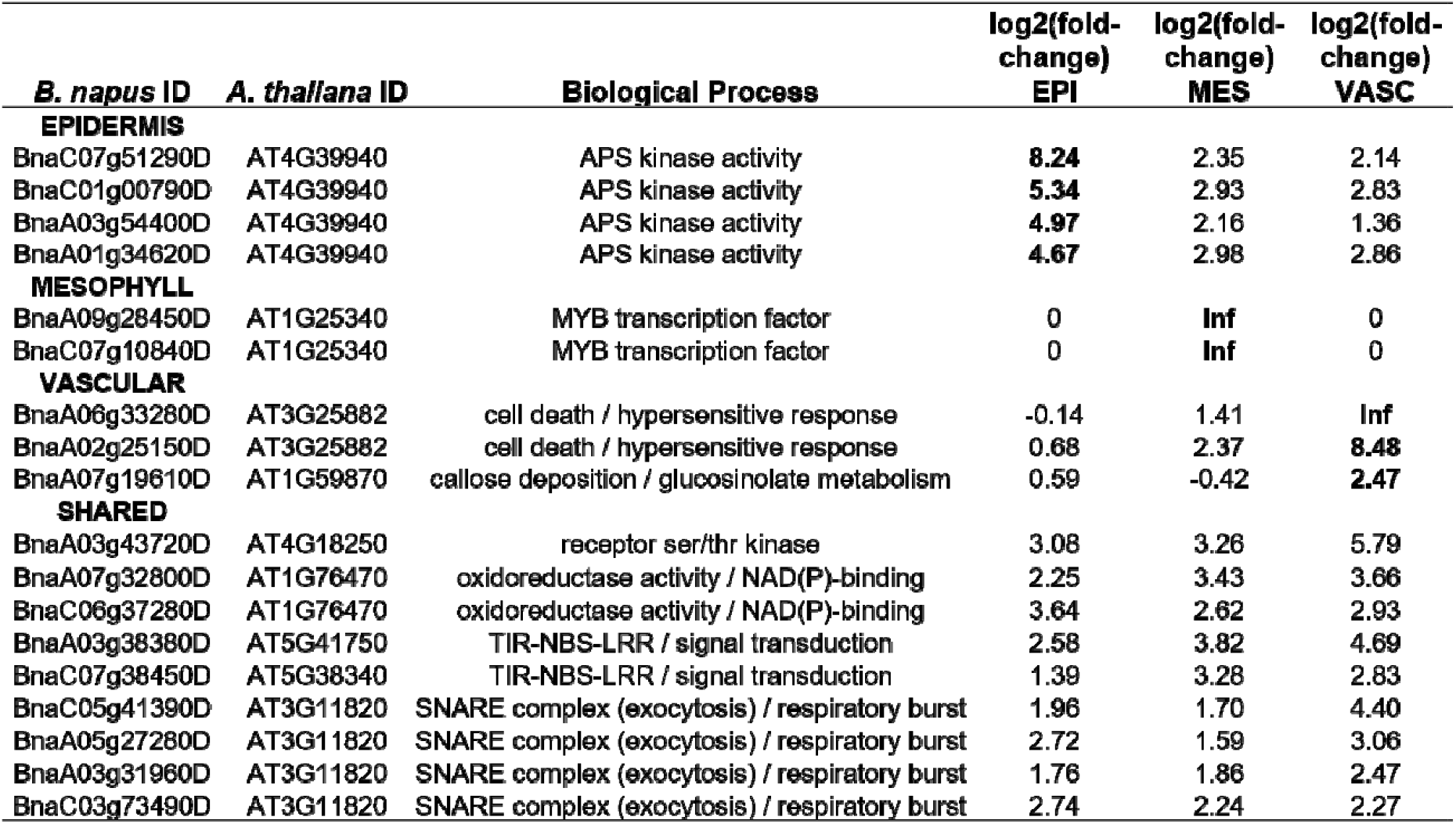
Abundance of tissue - specific and shared defense-related genes in response to *Scierotinia Sclerotiorum* infection Log2 fold - change represents degree of up regulation in *B. napus* tissue layers in response to infection. Candidate genes selected for functional charecterization using Arabidopsis.T-DNA Insertion lines in response to pathogen attack.

| *B. napus* ID | *A. thaliana* ID | Biological Process | log2(fold-change)<br>EPI | log2(fold-change)<br>MES | log2(fold-change)<br>VASC |
| --- | --- | --- | --- | --- | --- |
| **EPIDERMIS** |  |  |  |  |  |
| BnaC07g51290D | AT4G39940 | APS kinase activity | **8.24** | 2.35 | 2.14 |
| BnaC01g00790D | AT4G39940 | APS kinase activity | **5.34** | 2.93 | 2.83 |
| BnaA03g54400D | AT4G39940 | APS kinase activity | **4.97** | 2.16 | 1.36 |
| BnaA01g34620D | AT4G39940 | APS kinase activity | **4.67** | 2.98 | 2.86 |
| **MESOPHYLL** |  |  |  |  |  |
| BnaA09g28450D | AT1G25340 | MYB transcription factor | 0 | **Inf** | 0 |
| BnaC07g10840D | AT1G25340 | MYB transcription factor | 0 | **Inf** | 0 |
| **VASCULAR** |  |  |  |  |  |
| BnaA06g33280D | AT3G25882 | cell death / hypersensitive response | -0.14 | 1.41 | **Inf** |
| BnaA02g25150D | AT3G25882 | cell death / hypersensitive response | 0.68 | 2.37 | **8.48** |
| BnaA07g19610D | AT1G59870 | callose deposition / glucosinolate metabolism | 0.59 | -0.42 | **2.47** |
| **SHARED** |  |  |  |  |  |
| BnaA03g43720D | AT4G18250 | receptor ser/thr kinase | 3.08 | 3.26 | 5.79 |
| BnaA07g32800D | AT1G76470 | oxidoreductase activity / NAD(P)-binding | 2.25 | 3.43 | 3.66 |
| BnaC06g37280D | AT1G76470 | oxidoreductase activity / NAD(P)-binding | 3.64 | 2.62 | 2.93 |
| BnaA03g38380D | AT5G41750 | TIR-NBS-LRR / signal transduction | 2.58 | 3.82 | 4.69 |
| BnaC07g38450D | AT5G38340 | TIR-NBS-LRR / signal transduction | 1.39 | 3.28 | 2.83 |
| BnaC05g41390D | AT3G11820 | SNARE complex (exocytosis) / respiratory burst | 1.96 | 1.70 | 4.40 |
| BnaA05g27280D | AT3G11820 | SNARE complex (exocytosis) / respiratory burst | 2.72 | 1.59 | 3.06 |
| BnaA03g31960D | AT3G11820 | SNARE complex (exocytosis) / respiratory burst | 1.76 | 1.86 | 2.47 |
| BnaC03g73490D | AT3G11820 | SNARE complex (exocytosis) / respiratory burst | 2.74 | 2.24 | 2.27 |

First, we quantified lesion progression and fungal load relative to wild type Col-0 at 3 dpi with the necrotrophs *S. sclerotiorum* and *B. cinerea* (Fig. 3). Four of the five mutant lines that demonstrated a significant increase in lesion progression and fungal load relative to Col-0 were from the shared gene grouping (Fig. 3). The T-DNA insertion lines with increased susceptibility included *at4g18250* (receptor ser/thr kinase, *pr5k*), *at5g41750* (*tir-nbs-lrr* class protein), *at3g11820* (*pen1*), and *at1g76470* (*NAD(P)-binding protein*) from the shared group, and the epidermis-specific *at4g39940* (*apk2*). T-DNA insertion lines *pr5k* and *at5g41750*, which are putatively involved in PTI and ETI pathogen recognition respectively, had the strongest susceptibility phenotypes with a 310% and 221% increase in *S. sclerotiorum* fungal load respectively, and a 145% and 138% increase in *B. cinerea* fungal load relative to inoculated Col-0 plants.

**Figure 3.**
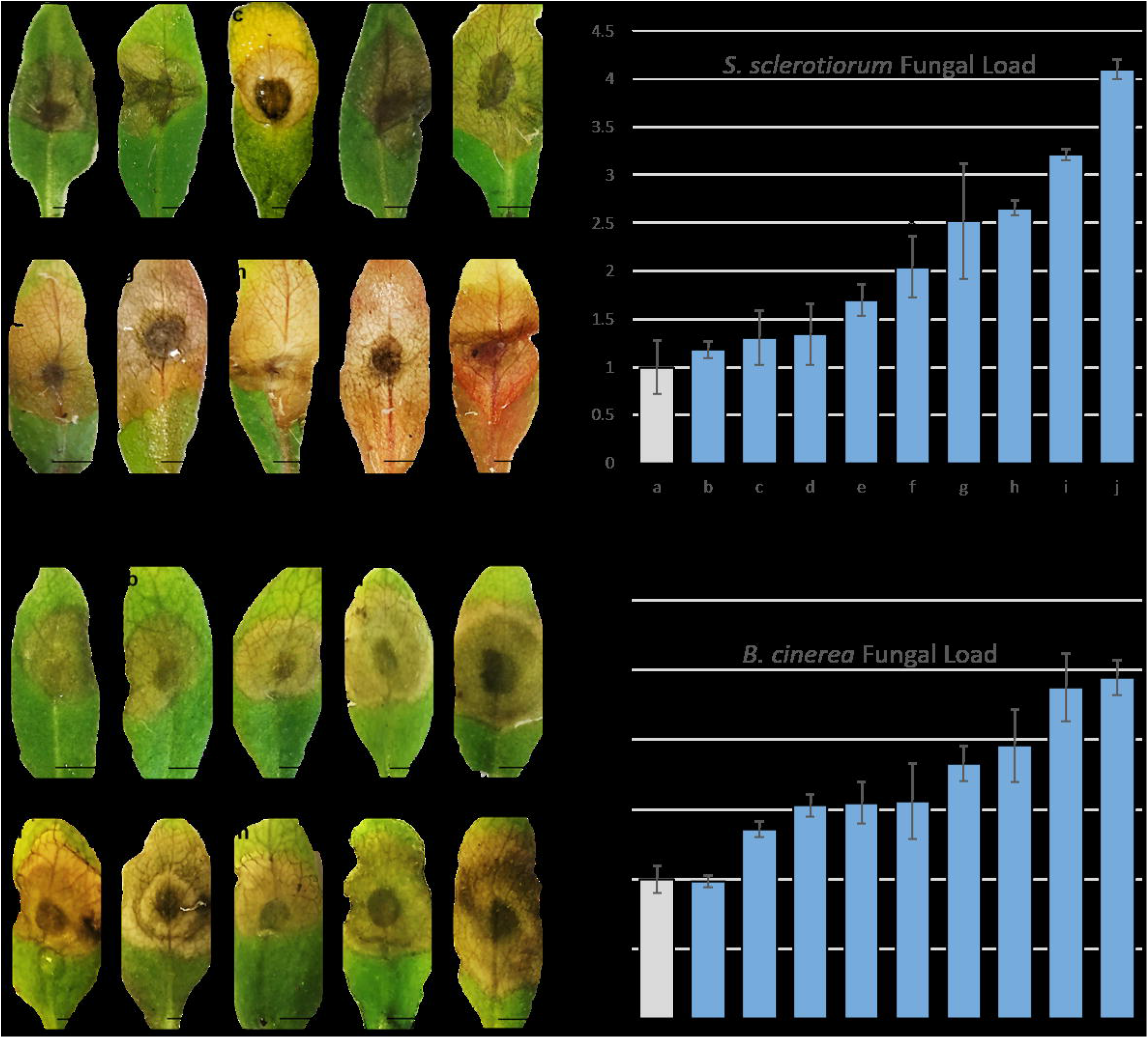
A) *S. sclerotiorium* and C) *B. cinerea* lesion development in wild-type Col-0 and T-DNA insertion Arabidopsis. Fungal load quantified using qPCR and relative transcript abundance against Col-0 at 3 dpi with ascospore PDB + Peptone solution for B) *S. sclerotiorum* and D) *B. cinerea*. Labels a – j represent Arabidopsis lines challenged. Error bars represent standard error and scale = 5mm.

We then challenged Arabidopsis mutants with the hemi-biotrophic fungal pathogen *L. maculans* (Fig. 4). Arabidopsis is not a natural host of *L. maculans*; however, if the defense response is sufficiently compromised, infection can occur (Becker et al., 2017). T-DNA insertion lines with significant infection symptoms included *at4g18250 (pr5k), at1g59870 (pen3), at5g38340, at3g11820, at4g39940*, and *at3g25882* with *pr5k* showing the most significant susceptibility phenotype. When insertion lines were challenged with the *biotrophic pathogens, H. arabidopsidis and P. syringae*, similar results were found to that of the hemi-biotroph *L. maculans*, with both *pr5k* and *pen3* showing significantly increased fungal load and colony counts, respectively (Fig. 4). Of the two targets showing consistent susceptibility phenotypes, *pr5k* belonged to the shared expression and *pen3* was identified in the vascular-specific gene set. While infection assays revealed a susceptibility phenotype in *pen3* against all three biotrophic pathogens tested, *pr5k* showed susceptibility against all five pathogens, regardless of the pathogen type.

**Figure 4.**
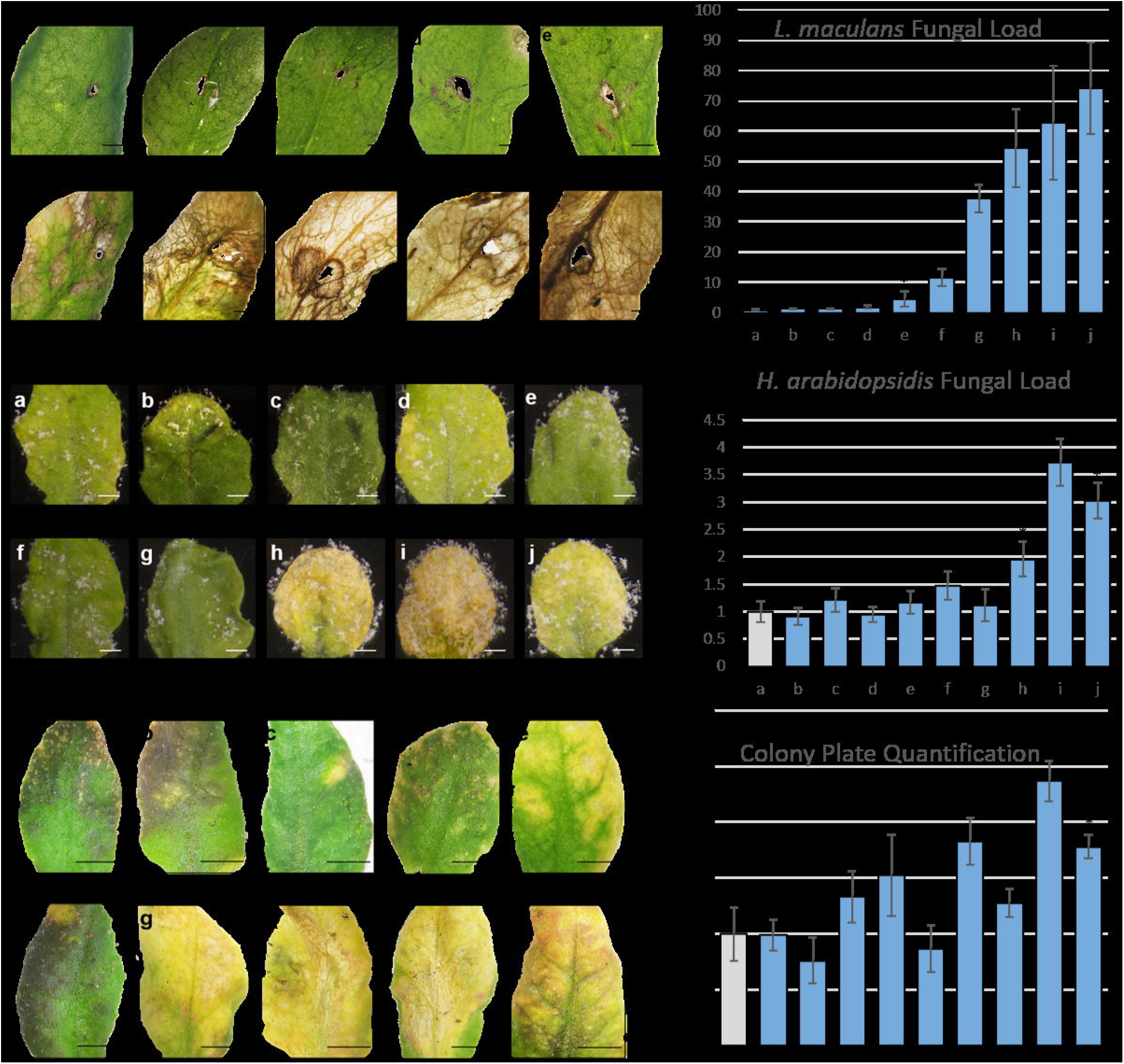
A) *L. maculans*, C) *H. arabidopsidis*, and E) *P. syringae* lesion development in wild-type Col-0 and T-DNA insertion Arabidopsis. Fungal load quantified using qPCR and relative transcript abundance against Col-0 at 20 dpi of D3 spore solution for B) *L. maculans, 10* dpi Emco5 spore solution for D) *H. arabidopsisidis*, and 5 dpi with DC3000 for F) *P. syringae. P. syringae* infected leaves were plated on Pseudomonas isolation media prior to colony quantification. Labels a – j represent Arabidopsis lines challenged. Error bars represent standard error, scale for A) = 1mm, and for B) and C) = 5mm.

### Dissection of PR5K receptor-like kinase signalling pathway in response to *S. sclerotiorum*

To better understand PR5K and its role in pathogen detection and downstream defense signalling, we quantified expression levels of predicted interacting partners using qPCR of wildtype Arabidopsis Col-0 and *pr5k* lines in response to *S. sclerotiorum* infection. Predictive interacting partners were identified using stringdb (https://string-db.org/) and co-expression data, and predictive transcription factor modules were identified in the FKM shared pattern using SeqEnrich (Becker et al., 2017). PR5K consists of an osmotin/thaumatin-like domain, a beta-1,3-glucanase domain at its N-terminal which may suggest potential activation through beta-1,3- glucan binding (Fig. 5), and a C-terminal serine/threonine protein kinase domain. Using stringdb and our global mRNA sequencing data, we identified four *B. napus* SnRK3s (SNF related kinases, subgroup 3) and CHOLINE KINASE 1 (CK1), which both demonstrate co-expression with *PR5K* in response to *S. sclerotiorum* infection (Fig. S1), suggesting a potential role in downstream signalling of PR5K. Using SeqEnrich to predict transcription factor gene regulatory modules, multiple transcription factors were identified to have the same co-expression pattern in response to infection and reduced expression in *pr5k* T-DNA insertion Arabidopsis, including LSH6 (AT1G07090) and four *B. napus* homologs of STZ (AT1G27730) transcription factors that were identified in our shared pattern. Our gene regulatory network analysis predicted LSH6 and the four STZ TFs to bind to the STZ DNA sequence motif upstream of genes belonging to GO terms response to chitin (P = 6.81×10^−52^), response to hypoxia (P = 4.34×10^−4^), and response to ABA (P = 2.44×10^−8^) (Fig. 5). Finally, we found a related but unknown RLK (AT5G48540) that showed significant up-regulation in the *pr5k* insertion line and potential functional conservation between the two RLKs (AT5G48540 and PR5K) (Fig. S1).

**Figure 5.**
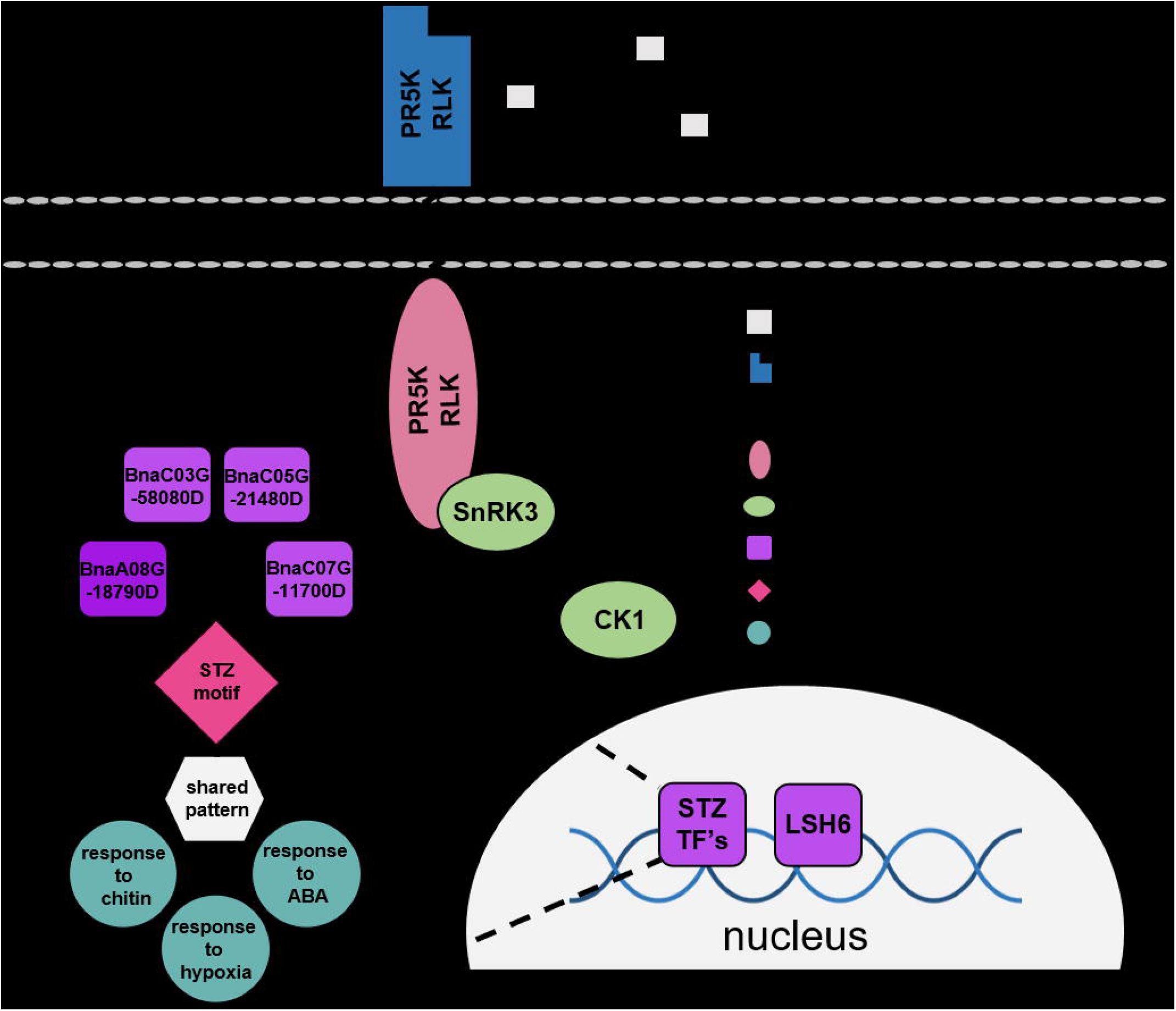
Predictive model of the PR5K receptor-like kinase (RLK) and its potential interacting partners. *S. Sclerotiorum* released pathogen associated molecular patterns (PAMPs) bind to the extracellular osmotin/thaumatin-like domain activating down-stream signal transduction on the intracellular serine/threonine kinase domain. Associated SNF-related kinase subgroup 3 (SnRK3) kinases are activated, potentially dependent on increased levels of ABA and can further act on down-stream kinases, such as choline kinase 1 (CK1). This signal transduction activates defense related transcriptional regulators like LSH6 and STZ transcription factors. STZ transcription factors can regulate expression of defense genes involved in the response to chitin, response to hypoxia, and response to ABA.

## Discussion

### *S. sclerotiorum* induces tissue-specific gene activity in the *B. napus* leaf

We coupled LMD and global mRNA sequencing to profile the three primary tissue systems of the *B. napus* leaf in response to *S. sclerotiorum* infection. As shown in previous studies (Chan et al., 2016, Becker et al., 2017, Ziegler et al., 2019), LMD increases the resolution of gene expression analyses by detecting lowly expressed tissue-specific transcripts which otherwise would not be detected using traditional whole-leaf or tissue sequencing. Here, LMD of the epidermis, mesophyll, and vasculature increased the number of transcripts identified in *B. napus* leaves by 29% or 19 300 transcripts compared to previous whole-leaf sequencing strategies (Girard et al, 2017). Clustering analysis of these data reveal distinct patterns of transcript abundance in vascular tissue compared to epidermis and mesophyll tissue. This is consistent with previous tissue-specific RNA-sequencing of funiculus tissue layers in *B. napus* (Chan et al., 2016) and may be attributed to the specialization of each tissue layer.

The epidermis is the first tissue layer to encounter the attacking pathogen and its secreted pathogenicity factors. At the onset of infection, *S. sclerotiorum* releases oxalic acid which is an essential pathogenicity factor that modulates the host-redox environment (Seifbarghi et al., 2017). Initially, oxalic acid suppresses the oxidative burst in host cells, resulting in a delayed production of host reactive oxygen species (ROS, Kim et al., 2008; Seifbarghi et al., 2017). Using FKM clustering, transcripts up-regulated specifically in the epidermis show a strong association with redox homeostasis, through the biosynthesis of polyamines, spermine and spermidine. Polyamines play a dual role in redox homeostasis. Spermine and spermidine have been shown to reduce oxidative stress in response to biotic and abiotic stressors (Chen et al., 2019; Gupta et al., 2016) through the activation of antioxidant enzymes leading to an increase in ROS scavenging (Tang and Newton, 2005). However, polyamines have also shown the ability to induce the host oxidative burst through the regulation of ROS synthesis (Saha et al., 2015). In support of these epidermis-specific polyamines inducing the oxidative burst, we find co-enrichment of ethylene-activated signalling. Ethylene has been shown to be a regulator of FLAGELLIN SENSITIVE 2 (FS2, BnaC09g19310D) which is required to initiate the host oxidative burst (Danna et al., 2011), and displays epidermis-specific up-regulation in response to infection. While it is possible that epidermis-specific polyamines may play a role in activating ROS scavengers to reduce damage from pathogen released ROS, our data suggests a role in polyamine associated activation of the host oxidative burst in the epidermal tissue layer.

In the mesophyll tissue layer, we find a strong enrichment of peroxidase activity coupled with the catabolism of the endogenous ROS, hydrogen peroxide. This suggests a response to increased ROS levels in mesophyll tissue once the oxidative burst has been initiated in the epidermis. In a prior study by Steiner-Lange et al., 2003, the peroxidase PR-9 accumulated specifically within the mesophyll tissue layer in response to *R. secalis* infection in barley leaves. PR-9 peroxidase activity is involved in cell wall reinforcement specifically through lignification, which is supported in our data through the mesophyll-specific enrichment of processes associated with cell wall modifications and reinforcement, and lignin biosynthesis. Further, peroxidases have been shown to form cross-links with glycoproteins to strengthen the cell wall in response to stress (Tenhaken, 2015). Ultimately, in response to pathogen attack, plant cell walls undergo a remodelling to reinforce physical barriers. Our data suggests this cell wall remodelling and reinforcement primarily occurs in the mesophyll to establish a strengthened physical barrier and obstruct the pathogen from reaching the vasculature tissue, thus preventing infection from becoming systemic.

The vascular tissue layer shows enrichment of induced resistance pathways, the JA and ET mediated ISR pathway, and the SA mediated SAR pathway in response to *S. sclerotiorum*. Multiple studies have described JA/ET signalling pathways as having strong up-regulation in response to necrotrophic pathogens like *S. sclerotiorum*, while SA is more commonly associated with host responses to biotrophic pathogens (Proietti et al., 2018; Li et al., 2019). To better understand this vascular-specific enrichment of SA and SAR in response to *S. sclerotiorum*, we examined expression levels of known molecular markers and key regulators of the SA and SAR pathways. PATHOGENESIS RELATED GENE 1 (PR1) and its positive regulator NON-INDUCIBLE IMMUNITY 1 (NIM1) are molecular markers of SA-mediated SAR (Weigel et al., 2005; Maier et al., 2011); however, both genes demonstrate down-regulation in the vascular tissue layer in response to *S. sclerotiorum*. This down-regulation is consistent with previous transcriptome analyses of *B. napus* responding to necrotrophic pathogens (Wu et al., 2016). Further, NIM-INTERACTING 1 (NIMIN1) is a negative regulator of NIM1 (Weigel et al., 2005) and demonstrates vascular-specific up-regulation in our dataset. Therefore, vascular-specific NIMIN1 activity is likely driving down-regulation of NIM1 and PR1 in vascular tissues of *B. napus*. While these data suggest inactivity of the SAR pathway in response to *S. sclerotiorum*, we detected strong vascular-specific up-regulation of the SA biosynthesis marker ISOCHORISMATE SYNTHASE 1 (ICS1). This increase in ICS1 suggests that SA could accumulate in the vascular tissue following Sclerotinia infection, but at 24 hpi, when the RNA samples were collected, levels of SA were insufficient to induce transcription of SAR-associated marker genes Indeed, SAR has previously been associated with later stages of pathogen infection taking multiple days to be active throughout the host (Hartman et al., 2016), and analysis of some later time points post-inoculation of *S. sclerotiorum* would help resolve whether this nectrophic fungus can promote an SA-induced SAR immune responses. Further, in support of vascular-specific ISR enrichment, we find ISR-involved JA and ET signalling pathways in vascular-specific gene co-expression patterns. While these data provide insight into vascular-specific SA and SAR activity in response to *S. sclerotiorum* infection, a time series through early, middle, and late stages of the infection process should provide additional insight into the complexity of tissue specific defense cascades in the plant.

### Pattern recognition and immunity pathways are shared between tissue layers of the leaf in response to *S. sclerotiorum*

In addition to tissue-specific patterns of gene activity, our co-expression analysis also identified shared patterns of gene activity in response to *S. sclerotiorum* with samples forming clusters based on response to infection. Within this shared co-expressed gene set, we identified biological processes associated with PTI and PRR signalling, coupled with serine/threonine kinase activity and down-stream signal transduction cascades (Tor et al., 2009). Due to the aggressive nature of *S. sclerotiorum* and its ability to secrete degradative enzymes that breakdown host tissue, all three tissue layers are likely to interact with PAMPs and therefore require expression of PRRs to elicit the necessary immune response. While our data reveal PRR genes like PR5K are expressed in all three tissue layers and at comparable levels, the role of each tissue layer on signal perception and transduction through these PRRs is still unknown. Wyrsch et al. (2014) showed expression of PRRs in multiple root tissue layers, however, the strength of the immune response varied across the layers, regardless of PRR expression levels, with a stronger response of PTI marker genes found in vascular root tissue compared to the epidermis. Homologous PTI marker genes in *B. napus* showed a similar trend of increased up-regulation in our dataset within the vascular tissue in response to infection, suggesting a potentially higher level of PAMP sensitivity in the vascular tissue layer. However, regardless of strength, co-expression of these processes indicates a shared role of PTI and associated down-stream stress response in all three tissue layers.

### Loss of shared candidate defense genes renders plants highly susceptible to pathogen attack

Several studies of plant responses to stress have suggested that genes with widespread expression across multiple tissues or stages of infection (i.e. shared genes) can have a major impact on protecting plants from pathogen attack (Gamboa-Tuz et al., 2018; Becker et al., 2017). Here, we were able to directly compare changes in expression of both shared and tissue-specific candidate defense genes, quantifying their responsiveness to different pathogens’ attack. Susceptibility screens showed that four of our shared candidates, most notably *pr5k, pen3, at5g41750 and at3g11820*, when mutated, rendered the plants more susceptible to fungal and bacterial pathogens than tissue-specific candidate defense genes. Thus, defense candidate genes with shared expression are more likely to enhance the defense response due to their conserved expression across tissue layers. Becker et al., (2017) demonstrated that genes showing conserved up-regulation in response to infection across multiple timepoints had greater impact on plant defense than candidate defense genes up-regulated in only one stage of infection. These findings combined with our tissue-specific analysis suggest gene candidates associated with plant defense that are expressed across both time and space are essential in guiding how effectively the plant responds to pathogen attack.

In addition to the differences observed between tissue-specific and shared gene candidates, we also uncovered host plant genes dependent on plants being challenged with either necrotrophic or biotrophic pathogens. The clearest example of this is *pen3*, whose mutants showed the strongest and most consistent susceptibility phenotype against biotrophic pathogens, but no such phenotype when challenged with necrotrophic pathogens. PEN3 is an ATP-binding cassette (ABC) transporter and is localized to the plasma membrane, playing a role in non-host resistance through the export of antimicrobial metabolites to the infection site (Stein et al., 2006; Campe et al., 2016). The lack of susceptibility of *pen3* mutants to necrotrophs may be due to PEN3 activity being dependent on its complexing partner PEN2, which together with PEN3 activates and subsequently exports key antimicrobial products including glucosinolates to the site of infection (He et al., 2019). While PEN3 does demonstrate up-regulation in vascular tissue in response to *S. sclerotiorum* infection, its complexing partner PEN2 is not co-expressed in our dataset. This suggests the PEN2/PEN3-dependent extracellular defense response may not be activated in response to necrotrophic pathogen attack and provides evidence into why *pen3* alone does not demonstrate a susceptibility phenotype against necrotrophic fungal pathogens. While *pen3* mutants showed significant and consistent susceptibility against biotrophs, our other top performing candidate *pr5k* mutants showed susceptibility phenotypes against all pathogens tested, regardless of mode of action. This consistent phenotype suggests PR5K activity is necessary for the host to mount a successful immune response against pathogen attack and is why the PRR served as a point of interest for down-stream analyses.

### PR5K acts as a central hub for pathogen detection and defense signal transduction essential for successful defense against pathogen attack

While previous studies have generated predictive models for the activation of the receptor serine/threonine kinase PR5K (AT4G18250) in response to abiotic stressors (Baek et al., 2019), activation of this receptor in response to biotic stressors like fungal and bacterial pathogens has yet to be explored. Fungal pathogens release pathogen associated molecular patterns (PAMPs), which are small molecular motifs capable of binding pathogen recognition receptors (PRRs) and initiating the plant’s immune response (Franco-Orozco et al., 2017; Zipfel et al., 2008). β1,3-D glucan has been shown to be secreted by fungal and bacterial pathogens as an exo-polysaccharide and can act as a PAMP to trigger immune responses through PRR binding (Batbayar et al., 2012; Oliveira-Garcia and Deising, 2013). PR5K contains a beta-1,3-glucanase domain within its extracellular osmotin/thaumatin-like domain, suggesting a potential interaction between the PAMP β1,3-D glucan and PR5K to initiate down-stream signalling.

PR5K also contains a transmembrane and an intracellular serine/threonine kinase domain. Previous studies have shown an association between the PR5K kinase domain and SnRKs which can be activated in response to intracellular abscisic acid (Baek et al., 2019). SnRKs are divided into three subgroups based on their modes of activation, subgroup 1 being ABA-independently activated, subgroup 2 being weakly activated by ABA, and subgroup 3 being ABA-dependent (Wang et al., 2020). We identified four *B. napus* subgroup 3 SnRKs (SnRK3s) co-expressed with PR5K in response to infection. This, coupled with the enrichment of response to ABA (P = 3.3×10^−6^) in our shared gene set and lack of co-expression of SnRK1 and SnRK2 subgroupings with PR5K, suggests a role of ABA in signal activation not only in response to abiotic stress, but also in response to *S. sclerotiorum* infection. The predicted interacting partner CHOLINE KINASE 1 (CK1) is also co-expressed with PR5K and shows a significant reduction in expression in *pr5k* Arabidopsis, providing additional evidence for its role in down-stream signalling. Transcription factor network analysis identified the transcriptional regulator LSH6, and four *B. napus* STZ transcription factors (BnaA08g18790D, BnaC03g58080D, BnaC05g21480D, and BnaC07g11700D) which showed significantly reduced expression levels in *pr5k* in response to infection and enrichment of host response to chitin, hypoxia, and ABA, further supporting the idea of an ABA-dependent activation of PR5K signal transduction as STZ activity has been shown to be induced in response to ABA (Sakamoto et al., 2004). Functional characterization of STZ transcription factor activity is required to identify its role in the enrichment of plant immunity processes. Lastly, the unknown receptor-like kinase (unk-RLK) (BnaC07g11700D) also demonstrates co-expression with PR5K, however, transcript abundance is increased in the *prk5* mutant (Fig. S1), suggesting a potential conservation of function with PR5K leading to unk-RLK up-regulation when PR5K is inactive. While these data provide insight into the activation and down-stream signalling activity of PR5K in response to infection, further characterization through sequencing of *pr5k* Arabidopsis post infection and analysis of down-stream protein-protein interactions are required to better understand this plant immunity pathway.

Taken together, our data provide insight into the role of tissue-specific and shared transcripts in the *B. napus* leaf in response to *S. sclerotiorum* infection. While several features of the defense response were shared across tissue layers, each tissue of the leaf accumulated specific mRNA populations in response to fungal infection. Despite tissue-specific activation of gene activity in response to pathogen attack, candidate shared defense genes out-performed their tissue-specific counterparts when challenged with biotrophic, hemi-biotrophic, and necrotrophic pathogens. Finally, our data suggests that the PRR PR5K plays a central role in successful host plant response to both fungal and bacterial pathogen attack through pathogen detection and activation of signalling cascades resulting in the enrichment of host immunity.

## Supplementary data

**Figure S1**. Predictive interacting partners with PR5K in response to *S. sclerotiorum*.

**Dataset S1**. *Brassica napus* FPKM abundance values.

**Dataset S2**. Pattern gene lists for fuzzy k-means clustering analysis.

**Dataset S3**. Gene ontology enrichment for fuzzy k-means clustering patterns.

**Dataset S4**. Fungal load and predictive PR5K partner qPCR primers.

**Dataset S5**. Transcripts detected between samples.

**Dataset S6**. *Brassica napus* – Arabidopsis homologous gene list.

## Acknowledgements

The authors would like to thank Dr. Michael Becker for his expertise in large scale data analysis, Dr. Sanjay Saikia for his help in the collection of leaf tissue through laser microdissection and the Genome Quebec CES facility for processing cDNA libraries and providing raw mRNA sequencing data. This work was supported in part through a Manitoba Agriculture Rural Development Initiatives Growing Forward 2 and a Canola Agronomic Research Program (CARP) grant through Canola Council of Canada to MFB, TDK, and WGDF. Additional funding was provided by the Natural Sciences and Engineering Research Council of Canada to MFB and SW.

## Author Contributions

PLW, IJG, MFB, SW, TRDK, and WGDF conceptualized and designed the study. PLW, IJG, SG performed experiments, and PLW performed data analysis. PLW wrote the manuscript. MFB, SW, WGDF, and TRDK supervised the study and contributed to the editing of the manuscript.

## Data availability statement

All RNA sequencing data are available at the Gene Expression Omnibus (GEO) data repository (GSE169299).

## Figure legends

**Figure S1. Predictive interacting partners with PR5K in response to *S. sclerotiorum* infection**. A) Co-expression table of target predictive interacting partners from shared up-regulation in response to infection pattern. B) qPCR target transcript abundance of predictive interacting partners in col-0 and *at4g18250* knockout Arabidopsis.

